# The effect of acute and prolonged acidosis on a panel of carcinoma cell lines

**DOI:** 10.1101/2025.06.19.660541

**Authors:** Maria Dravecka, Claire Wells, Ole Morten Seternes, Jakob Mejlvang

## Abstract

Due to cancer cell metabolism and disorganized tissue structure, the tumour microenvironment is associated with several pathophysiological conditions, including hypoxia, nutrient deprivation, accumulation of waste products, and acidification of the tumour microenvironment (tumour acidosis). Despite the belief that tumour acidosis drives tumorigenesis, it is still not clear how cancer cells respond to acidosis in the absence of other pathophysiological conditions. Here, we investigate how both acute and prolonged acidosis (pH 6.8) affects different epithelial features of a panel of carcinoma cell lines. We find that acute acidosis in all cell lines investigated represses the cell growth and causes a disturbance of adherens junctions and apical-basal polarity, reminiscent of epithelial-to-mesenchymal transition (EMT). However, we do not find that acute acidosis has a general effect on adhesion and migration in our panel of cell lines. Exposing our panel of carcinoma cell lines to acidosis for more than six weeks did not lead to adaptations restoring cell growth. On the contrary, prolonged acidosis caused one cell line (A431) to halt proliferation. Another cell line (A549) reacted to prolonged acidosis by gradually inducing the expression of ZEB2, which caused a partial EMT. In the rest of the cell lines, we did not find any noticeable effect of prolonged acidosis. Lastly, all cell lines quickly restored their original phenotype and growth rate when returned to media with normal pH (pH 7.5).

## Introduction

In humans, both blood and interstitial fluids have a slightly basic pH of approximately 7.3–7.5. Under non-pathological conditions, cells prefer oxidative metabolism, resulting in the conversion of fatty acids, amino acids, and carbohydrates into energy and the waste product CO_2_. This constant production of CO_2_ is neutralized to prevent a toxic acidification of the interstitial fluids within the tissue. This is facilitated by the bicarbonate buffer system, which enables the safe transport of CO_2_ via blood to the respiratory system. Solid tumours are highly heterogeneous structures, with distinct tissue organization and microenvironmental features from the core to the periphery. In denser regions of the tumour, diminished vasculature suppresses the flux of interstitial fluids and thereby reduces perfusion of the tumour tissue^1^. Proliferating cancer cells have comprehensively reprogrammed their metabolism and largely meet their energy needs through aerobic glycolysis, which leads to increased production of secretory lactate^2,3^. Due to these circumstances, both CO_2_ and lactic acid accumulate in the tumour tissue causing the extracellular pH of the tumour microenvironment to reach 6.5-7.0^4^. While the cytosolic pH of cells in this environment is generally maintained at a relatively stable level, acidosis still induces a slight reduction in intracellular pH^5,6^. However, the reduction in extracellular pH exceeds the reduction in cytosolic pH, which causes a reversed pH gradient across the plasma membrane^7,8^.

Cancer cells are directly affected by acidosis through both extracellular and intracellular mechanisms^2,9^. Enzymatic modulation of the extracellular environment is highly affected by pH, and acidosis could contribute to tumorigenesis by e.g. enhancing the degradation of the extracellular matrix^10^. Acidosis has also been shown to affect several intracellular mechanisms, including autophagy^11^ and cell metabolism^12–16^. For instance, cancer cells exposed to acidosis shift their preference from glucose to glutamine metabolism^12,13^ and increase fatty acid oxidation, which then becomes the main source of acetyl-CoA to support the TCA cycle^13,16^. However, acidosis has also been reported to induce cellular programs. For instance, acidosis can induce changes in morphology reminiscent of epithelial-to-mesenchymal transition (EMT)^17–25^. EMT is a cellular program that causes epithelial cells to lose their cell polarity, diminish adhesion to neighbouring cells, and gain migratory and invasive properties. EMT is induced by EMT-activating transcription factors, including ZEB1, ZEB2, TWIST, SNAIL1, and SLUG (SNAIL2), which all share the ability to directly repress epithelial genes (e.g., CDH1 coding for E-cadherin) and simultaneously activate genes associated with a mesenchymal phenotype (e.g., CDH2 coding for N-cadherin)^26^.

Collectively these observations have instigated the idea that acidification of the tumour tissue through various cellular mechanisms contributes to accelerated tumour progression. In conjunction with the observations that acidosis also influences angiogenesis^27^ and the capacity of immune cells to target and destroy cancer cells^28^, cancer therapies that exploit acidosis-associated effects on cells within the tumour microenvironment potentially represent a promising new frontier in oncology. The understanding of how cancer cells respond to acidosis is naturally a crucial prerequisite for developing targeted cancer therapies based on acidosis. Although several excellent research groups have investigated the effect of acidosis on different cancer cell lines, it remains an open question how cancer cells in general respond to acidosis and whether specific intracellular signalling pathways are triggered by acidosis^2,29^.

Intrigued by the possibility that EMT could be a general response to acidosis in carcinoma cells, we decided to investigate how a panel of carcinoma cell lines was affected by acute and prolonged acidosis.

## Results

### Cell growth is reduced during acute acidosis

To ensure the best comparability with existing literature, we selected a pH of 6.8 to represent acidosis—a widely utilized value in *in vitro* acidosis studies and an average representation of reported pH measurements of tumour interstitial fluids. As a strategic approach, we reduced the sodium bicarbonate content in the growth media to decrease the pH to 6.8 in media acclimatised to 37°C and 5% CO_2_. To maintain pH stability and mitigate fluctuations caused by cellular metabolism, we decided to refresh the growth media every day. The experiments were conducted on a panel of human cancer cell lines, including A431 (Epidermoid carcinoma), A549 (Non-small cell lung carcinoma), HCT116 (Colorectal carcinoma), HT29 (Colorectal carcinoma), and MDA-MB-231 (Breast carcinoma). These cancer cell lines were chosen because they originate from different types of carcinomas and because the effect of acidosis has previously been studied in these cell lines.

To elucidate how acute acidosis generally affects cancer cells in the tumour microenvironment, we first examined the impact of acute acidosis on cell growth. Using the CellTiter-Glo® assay to estimate the number of cells based on metabolic activity (intracellular ATP), we observed a significant and similar reduction in cell growth across all five cell lines (Figure 1). The inhibitory effect of acidosis on cell growth was evident as early as 24 hours after induction (Figure 1) and caused a reduction ranging from 47% to 70% after three days, among the five cell lines (Figure 1 and Source Data file). Similar results were obtained through manual cell counting (Supplementary figure 1). In conclusion, all investigated cell lines exposed to acute acidosis continued to grow; however, with a slightly reduced growth rate. We did not observe any signs of increased cell death such as cellular debris. This indicates that acute acidosis causes a consistent and moderate suppression of cell growth across all tested cell lines.

**Figure 1.**
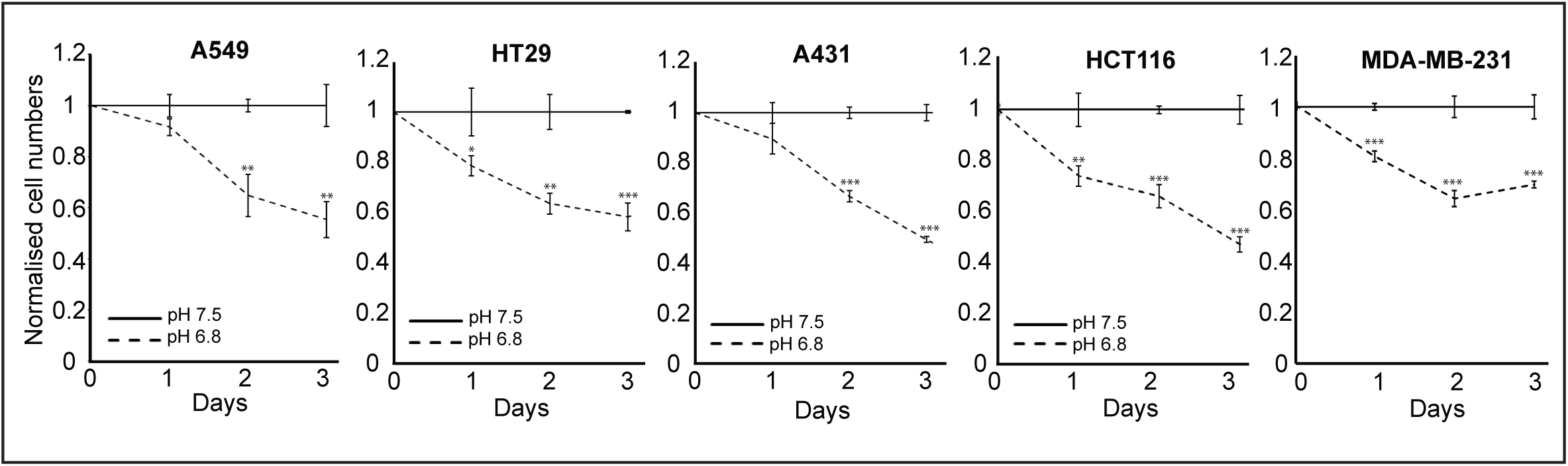
Low extracellular pH reduces the growth of cancer cells. Depicted cell lines were cultured for 3 days in media with either pH 7.5 or 6.8. Cell numbers (based on ATP levels) were measured daily using the CellTiter-Glo assay. The graphs illustrate results standardized to the cells cultured in media with pH 7.5. Error bars represent the standard deviation based on three technical replicates. The experiment was performed three times giving similar results. Statistical analysis was performed using two-tailed Students t-test; *p<0.05, **p<0.01, ***p<0.001. Source data are provided in the Source Data file.

### Acute acidosis perturbs adherens junctions and apical-basal polarity

The morphology of A431, HT29 and HCT116 cells is commonly referred to as cobblestone morphology, as these cells form dense epithelial sheets with well-defined intercellular adherens junctions. When these cells were exposed to acidosis, we observed a disruption of the well-organized cobblestone morphology within 24 hours, manifested by less distinct intercellular borders (Figure 2A). In the HCT116 cell line, we also noted a tendency of the epithelial sheet to scatter into individual cells with reduced contact with neighbouring cells (Figure 2A). Additionally, these cells displayed a tendency to flatten out, which could indicate a reduction of the apical-basal polarity. Although A549 and MDA-MB-231 cells are epithelial cancer cell lines, they do not display the classical cobblestone morphology. The only observed change in morphology caused by acute acidosis in these cell lines was increased cell flattering (Figure 2A and Supplementary figure 2A-B). To further elucidate the effects of acidosis on epithelial morphology in A431, HCT116 and HT29 cells, we decided to investigate the potential effect of acute acidosis on the adherence junctions and the focal adhesion. To this end, cells were stained for E-cadherin and Paxillin and analysed by fluorescence microscopy. In cells cultured in media with pH 7.5, E-cadherin localized distinctively in adherens junctions and exhibited a robust cobblestone-like pattern (Figure 2B). After two days of exposure to acidosis, the E-cadherin localization had become less distinct although it was still mainly localized to the intracellular junctions between cells. Z-stack analysis revealed that E-cadherin junctions were distributed in long distinct lateral stretches, perpendicularly to the basal membrane, running from the apical to the basal membrane in cells cultured at pH 7.5 (Figure 2C). In cells exposed to acute acidosis, these lateral stretches became shorter, less distinct, and less perpendicular to the basal membrane (Figure 2B, compare red boxes). Analysis of Paxillin and F-Actin at the basal membrane revealed no obvious impact of acute acidosis on localization and presence of focal adhesions (Figure 2C and Supplementary figure 3). These observations clearly demonstrate that acute acidosis causes a weakening of adherens junctions and a partial collapse of apical-basal polarity in epithelial cancer cells.

**Figure 2.**
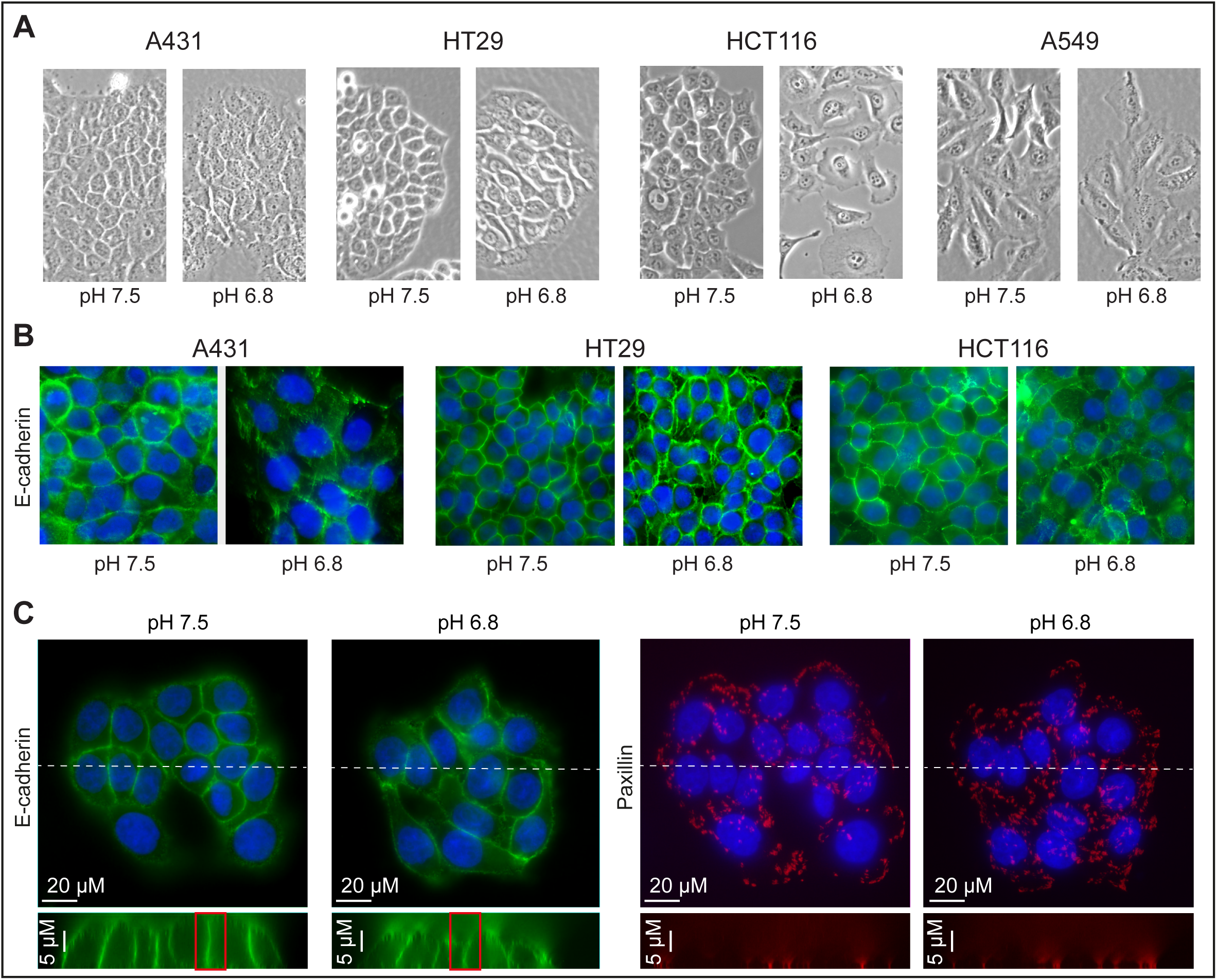
Low extracellular pH affects intercellular junctions. Cells were cultured in growth media with either pH 7.5 or pH 6.8 for two days. **A**, Images of cell morphology obtained by phase-contrast microscopy. **B**, Immunofluorescence micrographs depicting E-cadherin staining (green) in A431, HCT116, and HT29 cells. Blue depicts nuclei stained with DAPI. **C**, Immunofluorescence micrographs depicting E-Cadherin (green) and Paxillin (red) staining in HT29 cells. The bottom panel shows the localisation of E-cadherin and Paxillin through an orthogonal view at the transect indicated by the white dotted line.

### Acute acidosis does not affect E-cadherin expression

We next investigated whether acute acidosis induced cadherin-switching^30,31^ (downregulation of E-cadherin concomitant with upregulation of N-cadherin) and determined whether the relative expression levels of E- and N-cadherin were affected by acute acidosis. To this end, we harvested both RNA and protein from A549, A431, HCT116, and HT-29 cells before and 48 hours after exposure to acidosis. Of note, MDA-MB-231 does not express a noticeable amount of E-cadherin^32^. As a positive control, we also harvested protein and RNA from A549 cells stimulated with transforming growth factor β (TGF-β) for 48 hours, which has previously been shown to induce EMT including cadherin switching^33,34^. Western blot analysis confirmed that all cell lines expressed E-cadherin; however, the expression was not affected by acute acidosis. Intriguingly, we detected a small increase in RNA coding for E-cadherin in all cell lines exposed to acute acidosis (Figure 3B and C). In contrast, TGF-β stimulation of A549 cells caused a strong downregulation of E-cadherin at both the mRNA level (CDH1) and the protein level (Figure 3A and 3B). We could not detect any expression of the N-cadherin protein in HCT116, A431, and HT29 cells regardless of the pH of the growth media (Figure 3A), and nor did we detect any mRNA coding for N-cadherin (CDH2) by qPCR. In A549 cells, which are known to express modest level of N-cadherin under normal growth conditions^35^, we detected N-cadherin at both the protein and RNA levels (Figure 3A and B). Apparently, acute acidosis caused a small downregulation of N-cadherin at the protein level, while the expression of mRNA coding for N-cadherin (CDH2) did not change significantly (Figure 3B). In contrast, TGF-β stimulation of A549 cells caused a strong upregulation of N-cadherin on both mRNA and protein levels (Figure 3A and B).

**Figure 3.**
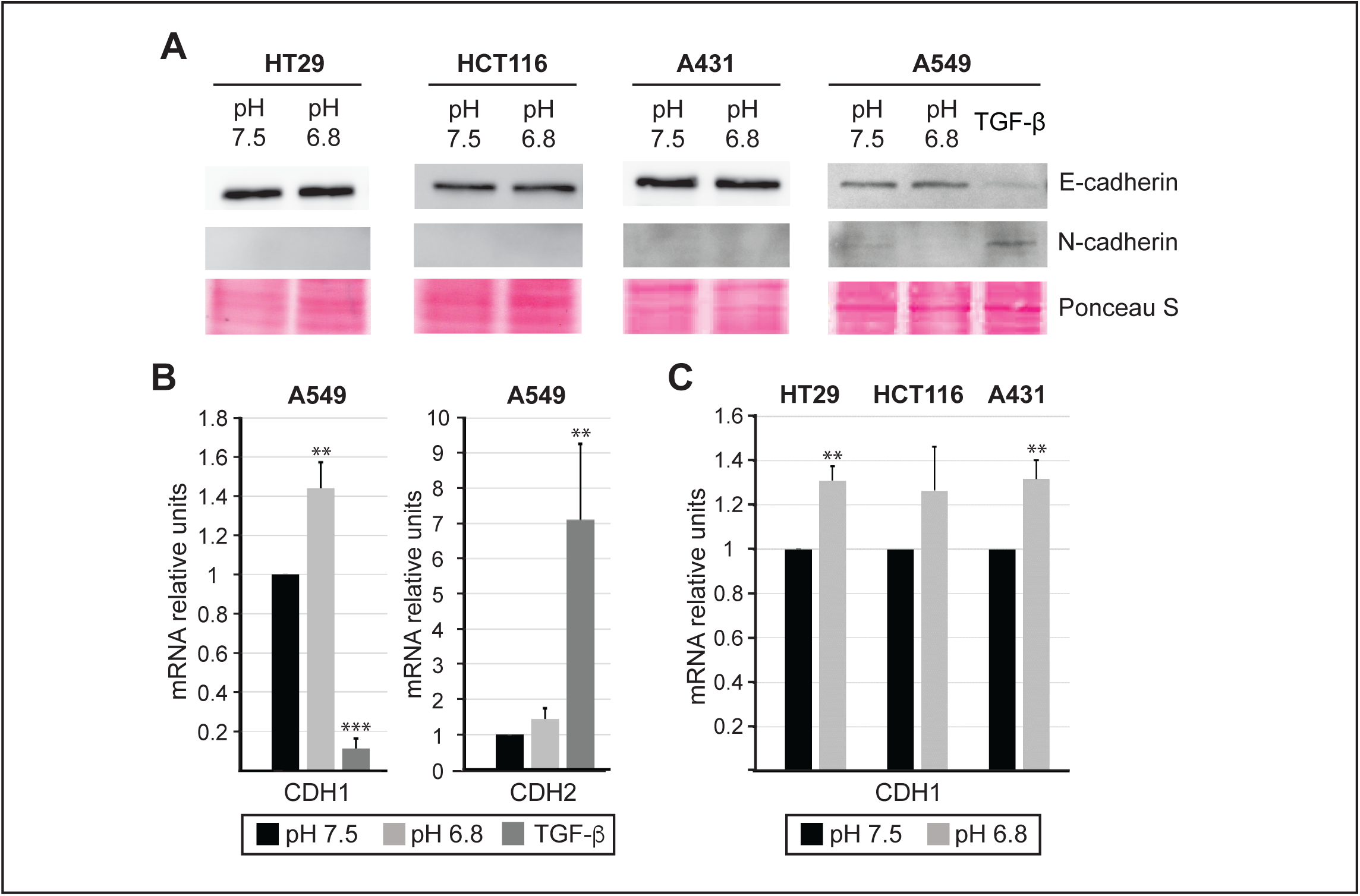
The expression of N-cadherin and E-cadherin is minimally affected by 2 days of acidosis. Indicated cell lines were cultured in media with either pH 7.5 or 6.8 for 2 days. Where indicated, A549 cells were exposed to media (pH 7.5) containing TGF-β (5 ng/ml) to serve as a positive control for EMT. **A**, Western blot analysis showing the protein expression of E-cadherin and N-cadherin. **B**, qPCR analysis revealing the relative expression of mRNA coding for E-cadherin (CDH1) and N-cadherin (CDH2) in A549 following the depicted treatments. The error bars represent the standard deviation of biological triplicates, and the results were standardized to the expression detected in cells cultured in media with pH 7.5. Statistical analysis was performed using two-tailed Student’s t-test; **p<0.01, ***p<0.001. Source data are provided in the Source Data file. **C**, qPCR analysis revealing the relative expression of mRNA coding for E-cadherin (CDH1) and N-cadherin (CDH2) in HT29, HCT116, and A431 cells cultured in media with either pH 7.5 or 6.8. The error bars represent the standard deviation of biological triplicates, and the results were standardized to the expression detected in cells cultured in media with pH 7.5. Statistical analysis was performed using a two-tailed Student’s t-test; **p<0.01. Source data are provided in the Source Data file.

In summary, we find that acute acidosis elicits morphological changes reminiscent of EMT, including disturbance of adherens junctions and partial loss of apical-basal polarity. However, these phenotypic changes are not coinciding with reduced expression of E-cadherin. Neither are they accompanied by an upregulation of N-cadherin.

### The effect of acute acidosis on adhesion and wound healing

The metastatic development of cancer cells necessitates a series of changes, including altered adhesion affinity to extracellular substrates and the acquisition of increased migratory and invasive potential. Therefore, we sought to investigate how acidosis affects these classical cell properties. First, cell adhesion was evaluated on two key extracellular matrix glycoproteins, namely Collagen and Fibronectin. Laminin was used as a negative control. All cell lines were cultured in media with either pH 6.8 or 7.5 for two days before performing the adhesion assays. In addition, we used A549 cells stimulated with TGF-β as a positive control for increased adhesive potential for Collagen and Fibronectin^34^. As expected, TGF-β-stimulated A549 cells displayed significantly increased adhesion to both Collagen and Fibronectin compared to control samples (Supplementary figure 4). Exposing HT29 and HCT116 cells to acidosis, did not affect adhesion to either Collagen or Fibronectin (Figure 4A). A549 cells subjected to acidosis exhibited decreased adhesion to Collagen, whereas adhesion to Fibronectin was not affected. The opposite effect was observed in A431 cells, where cells cultured under acidosis did not change their adhesion to Collagen but exhibited decreased adhesion to Fibronectin.

**Figure 4.**
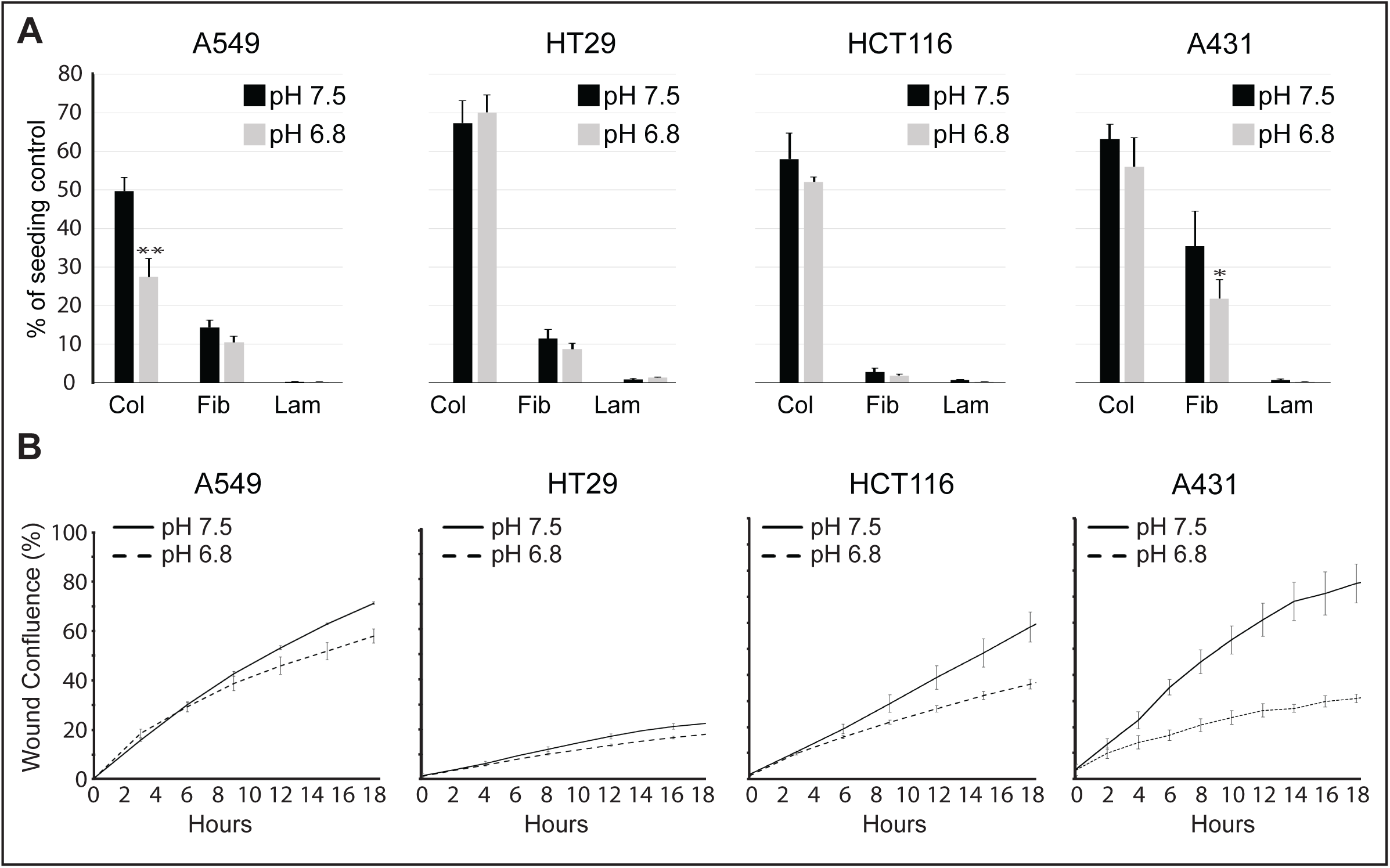
The impact of acidosis on adhesion and migration. Indicated cell lines were cultured in media with either pH 7.5 or 6.8 for 2 days. **A**, The relative adhesion to collagen (Col), fibronectin (Fib), and laminin (Lam) was assessed for each depicted cell line. Error bars represent the standard deviation of technical triplicates. The experiment was performed 3 times, yielding similar results. Statistical analysis was performed using two-tailed Student’s t-test; *p<0.05, **p<0.01. **B**, Wound healing assay was performed to assess cell migration. The error bars represent the standard error of technical replicates (n>5). The experiment was performed three times, yielding similar results. Representative images of wounds are shown in Supplementary figure 5.

Next, we tested whether acidosis affected wound healing, a fundamental process where epithelial cells proliferate and migrate to repair disrupted epithelium. Again, all cell lines were cultured under standard or acidosis conditions for two days prior to the experiment. Scratch wounds were meticulously created in confluent cell cultures, and wound closure dynamics were monitored and captured using the Incucyte® Live-Cell Analysis System. Acidosis treated A549, HT29, and HCT116 cells exhibited a slight tendency toward delayed wound closure (Figure 4B and Supplementary figure 5). However, since acidosis suppresses cell growth in these cells (Figure 1), we cannot exclude that the effect on wound closure is merely caused by the repressed cell growth. In contrast, acute acidosis clearly affected wound-closure in A431 cells. Cells cultured under standard conditions on average closed 75% of the wound within 18 hours, while those under acidosis only closed 31% of the wound (Figure 4B). Consistently, we observed that collective migration ceased in A431 cells when exposed to acidosis (data not shown).

Although we did observe some specific changes in adhesion and migration caused by acute acidosis, we conclude that acute acidosis does not elicit a general response affecting cell adhesion or migration in our panel of epithelial cancer cell lines.

### Prolonged effects of acidosis on epithelial cancer cells

Several reports have shown that cells can adapt to acidosis after prolonged (>6 weeks) exposure^12,19,25,36,37^. We therefore questioned whether we could detect any differences between acute and prolonged acidosis in terms of cell growth and morphology. To this end, we seeded all five cancer cell lines on tissue plastic and assessed the average daily growth rate within each passage for 6 consecutive weeks. During this time, the culture media was changed every second day, and the pH value of the waste media was measured and found to deviate by less than 0.2 from the expected 6.8. Cells exposed to acidosis for at least 6 weeks; were designated as “Acidosis experienced”. After two passages, A431 cells exposed to acidosis practically ceased to grow (Supplementary figure 6A). Due to the lack of growth, these cells no longer became confluent enough to be passaged and we could therefore not continue the assessment of the daily growth rate (see Materials and Methods). After six weeks, A431 cells were still sub-confluent (Supplementary figure 6C). However, when these cells were returned to media with normal pH, the growth rate slowly recovered (Supplementary figure 6B). In contrast, HCT116, HT29, MDA-MB-231, and A549 cells exposed to acidosis grew with a slightly reduced, but constant, growth rate throughout the entire six weeks (Figure 5A and Supplementary figure 7A). At the end of the experiment, when these acidosis experienced cells were returned to media with pH 7.5, the average daily growth rate returned to normal (Figure 5B and Supplementary figure 7B). Throughout the experiment, we consistently examined the morphology by phase contrast microscopy. Besides the already described morphological changes observed after two days of acidosis (Figure 2) we did not observe any effect of prolonged acidosis on cell morphology in HCT116 and HT29 (Figure 5C). In contrast, the morphology of A549 cells exposed to prolonged acidosis clearly caused individual cells to become more elongated and spindle shaped (Figure 5C and Supplementary figure 8A). In agreement, cell shape analysis of individual cells showed that both cell area and circularity, were significantly decreased in acidosis experienced A549 cells (Supplementary figure 8B). When acidosis experienced cells were returned to media with a pH of 7.5, their morphology reverted to the morphology of cells that had not experienced acidosis (Figure 5C and Supplementary figure 6C).

**Figure 5.**
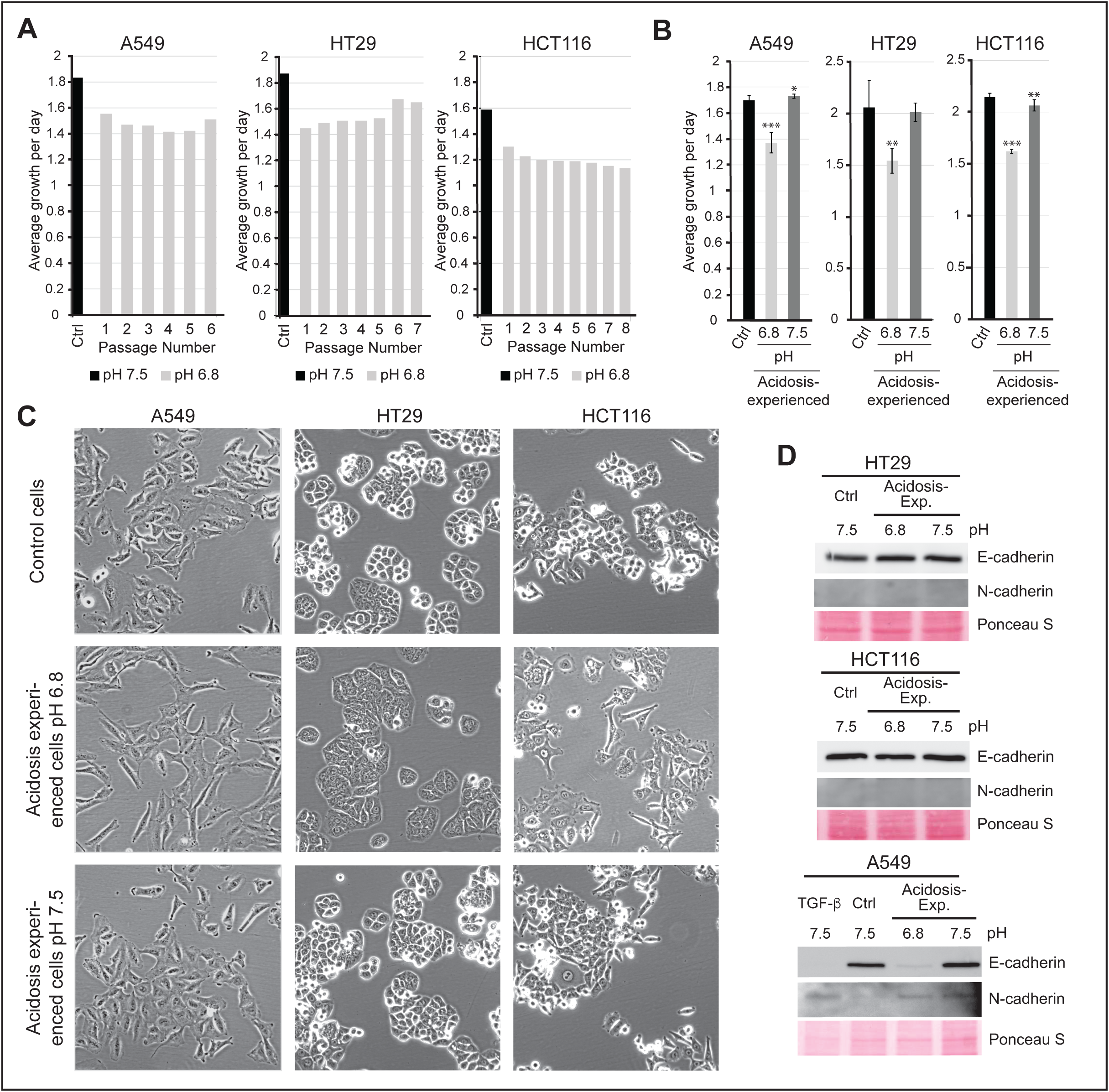
Prolonged acidosis can induce a reversible EMT. **A**, A549, HT29, and HCT116 cells were cultured for a minimum of 6 weeks in media with pH 6.8. Average daily growth is displayed for each consecutive passage following the start of the experiment. The experiment was repeated three times, yielding similar results. Source data are provided in the Source Data file. **B**-**D**, Cells cultured in media with pH 6.8 for more than six weeks (acidosis-experienced cells) were returned to media with pH 7.5 for one week. **B**, Average daily growth displayed for the depicted conditions. Error bars represent the standard error of technical triplicates. Statistical p-values were calculated using two-tailed Student’s t-test; *p<0.05, **p<0.01, ***p<0.001. Source data are provided in the Source Data file. **C**, Images of cell morphology obtained by phase-contrast microscopy. **D**, Western blot analysis of the relative expression of E- and N-cadherin.

In addition, we checked whether prolonged acidosis affected the protein expression of N- and E-cadherin. Whereas we did not detect any expressional changes in HT29 and HCT116 cells, we observed a significant downregulation of E-cadherin and upregulation of N-cadherin in A549 cells (Figure 5D). Similar to the effects on morphology and cell growth, these expressional changes were reverted when the acidosis experienced A549 cells were cultured in control media with pH 7.5. In general, we find that the observed effects of acute acidosis on cell growth and morphology are stable over time, and we did not observe any evidence suggesting that cells adapt to acidosis during the six weeks of exposure. However, one of the cell lines (A431) seemingly went into a dormant state associated with an extremely low growth rate. When reintroduced to media with pH 7.5 after the six weeks of acidosis, all cell lines restored their growth and morphology.

### Prolonged acidosis induces EMT in A549 cells

The observation that prolonged acidosis in A549 cells caused a strong decrease in E-cadherin and a moderate increase in N-cadherin intrigued us. We therefore questioned whether this was caused by transcriptional regulation. qPCR analysis revealed that prolonged acidosis significantly reduced the expression of CDH1 and increased the expression of CDH2 (Figure 6A). Similar to the protein levels (Figure 5D), the expression of these genes was restored when cells were reintroduced to media with pH 7.5 (Figure 6A). These findings substantiated that prolonged acidosis increased the expression of at least one of the transcriptional repressors of CDH1, namely SNAIL1, SNAIL2, ZEB1, ZEB2 and TWIST1. We therefore compared the relative expression of mRNA coding for these genes in A549 cells exposed to long-term acidosis to the level expressed in A549 cells cultured at normal pH (Figure 6B and Supplementary figure 9). Interestingly, long-term acidosis caused a significant upregulation of both ZEB2 and SNAIL2 (Figure 6B). This upregulation was completely reverted when acidosis experienced cells were reintroduced to media with pH 7.5 (Figure 6B), which coincided with the reversion of N- and E-cadherin protein expression (Figure 5D). To elucidate the temporal dynamics of acidosis-induced EMT in A549 cells, we analysed how the mRNA expression of CDH1, CDH2, SNAIL2 and ZEB2 was affected on a weekly basis after exposure to acidosis. This revealed that the relative expression of ZEB2 already increased within the first week of acidosis and continued to increase gradually over the following three weeks, after which it stabilized (Figure 6C). In contrast, the expression of SNAIL2 was found to fluctuate between 100% and 300% compared to control cells. The mRNA expression of CDH1 gradually decreased during the first 4 weeks, after which it stabilized at 15%. The opposite dynamics was observed for CDH2, which increased during the first 4 weeks and then it stabilized at approximately a 2.7-fold increase. In summary, this shows that one of our cell lines (A549) responds to acidosis by inducing a partial and reversible EMT, mediated mainly by ZEB2 but potentially assisted by SNAIL2.

**Figure 6.**
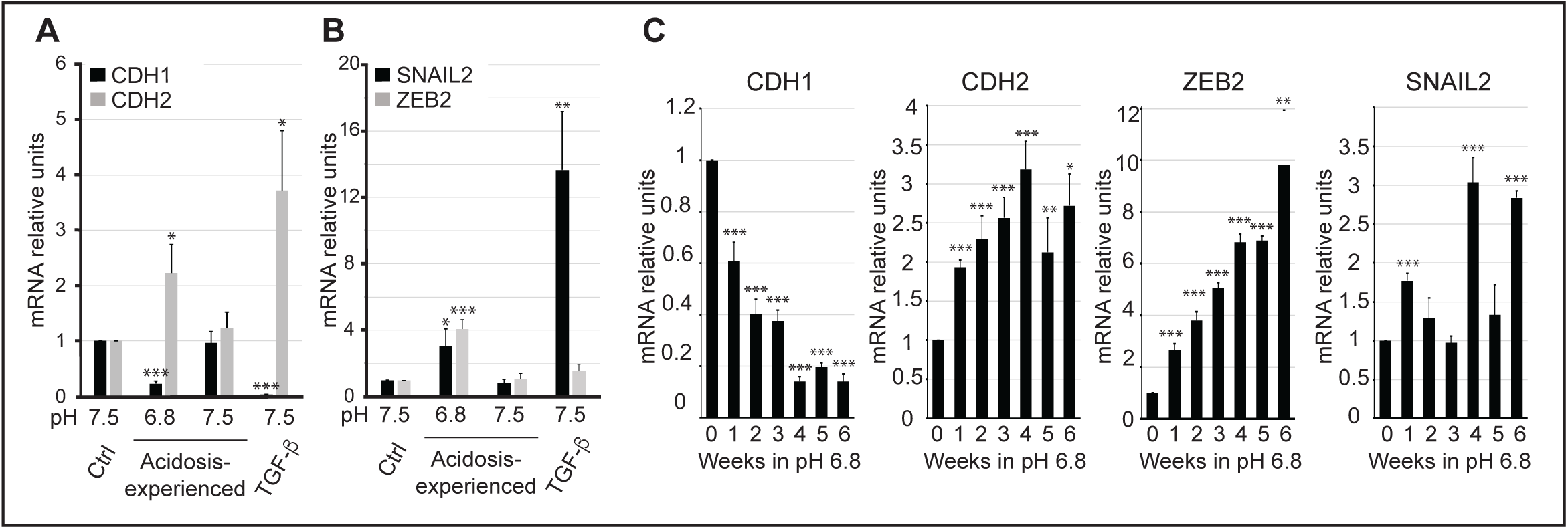
Prolonged acidosis gradually induces the expression of ZEB2 in A549 cells. A-B, The expression of mRNA coding for E-cadherin (CDH1), N-cadherin (CDH2), SNAIL2, and ZEB2 was assessed by qPCR in A549 cells following the depicted treatments. “Acidosis-experienced” refers to A549 cells that had been cultured for more than 6 weeks in media with pH 6.8. The TGF-β1 sample served as a positive control and derives from A549 cells cultured for 2 days in media (pH 7.5) containing 5 ng/ml TGF-β1. Error bars depict the standard deviation of technical triplicates. Statistical p-values were calculated using two-tailed Student’s t-test; *p<0.05, **p<0.01, ***p<0.001. Source data are provided in the Source Data file. **C**, The expression of the depicted genes was assessed weekly in A549 cells cultured in media with pH 6.8 for 6 weeks. Error bars depict the standard deviation of technical triplicates. Statistical p-values were calculated using two-tailed Student’s t-test; *p<0.05, **p<0.01, ***p<0.001. Source data are provided in the Source Data file.

## Discussion

The results of this study provide valuable insights into the effects of acute and prolonged acidosis on epithelial cancer cells, revealing both common and cell line-specific responses. One of the key challenges in understanding how acidosis impacts cancer cells lies in the variability of experimental setups and strategies used by various research groups. We demonstrated here that media with the pH of 6.8 causes a mild repression of growth in A549, A431, MDA-MB-231, HCT116 and HT29 cells, while it has previously been shown that the growth of HCT116 and HT29 cells ceases when cultured in media with pH 6.5^12,25^. Naturally this reflects that the effect of acidosis is dependent on its severity^21,38^. It is therefore important to stress that our investigations of how cancer cells are affected by acidosis are limited to acidosis induced at pH 6.8. Comparing the effect of acidosis (with the same severity) on cell growth between different cell lines demonstrated that cells can be differentially affected. There could be several reasons for this. Firstly, the cytosolic pH varies between different cancer cell lines^5,39^, and it is therefore likely that the severity of acidosis is directly linked to these variations. Secondly, the ability to maintain pH homeostasis as well as the metabolic production of acids are also varying between different cell lines^29^, and are therefore also potentially affecting the effect of acidosis.

Based on our observations we conclude that acute acidosis affects cancer cells of epithelial origin in a way that causes a slight reduction in their proliferation and a shift in epithelial morphology characterized by decreased intercellular adhesion and reduced apical-basal polarity. This is by no means controversial, given that acidosis has previously been reported to influence both proliferation and morphology^11–13,19,36,38,40^. As the morphological alterations were not accompanied by reduced expression of E-cadherin on either protein or RNA level, we conclude that these alterations are not caused by an induction of EMT *per se*. There is currently very little evidence suggesting that a conventional signal transduction pathway mediated by transcription is involved in eliciting a general response to acidosis. Firstly, because a transcriptional H^+^-responsive element has never been identified, and secondly because analysis of the transcriptomes from different cell lines exposed to acute acidosis has not led to the identification of genes ubiquitously regulated by acute acidosis^41,42^. However, it is possible that acidosis, rather than inducing a response, is causing an effect. Extracellularly, the decrease in pH increases the protonation of proteins and phospholipids embedded in the membrane which could directly affect their functioning^43–45^. Additionally, the activity and function of various extracellular enzymes, particularly those involved in proteolysis and extracellular matrix (ECM) degradation, are affected^46^. The minor cytosolic decrease in pH associated with acidosis^5^ also influences both transformed and non-transformed cells^47–50^. E.g. Phosphofructokinase-1, a rate-limiting key enzyme in glycolysis, is strongly inhibited when pH drop from 7.4 to 6.8^48^. Consistently acidosis was shown to reduce glycolysis in a large panel for cancer cell lines^47,49^. Decreased pH has also been shown to inhibit the Rho GTP-ases^50^ which are essential regulators of actin reorganization and consequently function in multiple cellular processes including cell migration and maintenance of apical-basal polarity in epithelial cells. These two examples show that even a small decrease in cytosolic pH can affect both proliferation and morphology. It is therefore not unlikely that the effect of acidosis on proliferation and morphology arises from a cumulative effect of myriads of subtle changes in molecular reactions caused by minor changes in pH. In contrast to proliferation and morphology, we did not observe that acidosis caused a general effect on adhesion or migration.

We did not see any evidence supporting the notion that epithelial cancer cells in general adapt to acidosis. However, we observed interesting cell-specific responses to prolonged acidosis. While four of the five cell lines investigated, maintained their slightly reduced growth rates during prolonged acidosis, A431 cells entered a dormant-like state with severely reduced proliferation. This has been observed in many other cell lines^25,51^, when the extracellular pH is lowered to 6.5. Perhaps that A431 cells are particularly sensitive to acidosis and therefore mimic other cells exposed to pH 6.5. This observation raises questions about the potential role of acidosis in tumour dormancy and warrants further investigation. In A549 cells, continuous acidosis repeatedly induced a gradual decrease in E-cadherin lasting for at least 4 weeks. As this coincided with increased expression of N-cadherin, ZEB2, and SNAIL2, we conclude that prolonged acidosis in A549 cells induces a partial EMT. We acknowledge that much more experimental work is needed in order to elucidate how acidosis mechanistically induces the observed partial EMT in A549 cells. However, since it is reversible and highly reproducible, we speculate whether this represents an adaptation to acidosis. In general, we find it challenging to distinguish between adaptation and transformation when it comes to cellular changes caused by long-term exposure to acidosis, especially when the chronological development is not analysed.

Our findings highlight the critical need for a deeper understanding of how acidosis influences both transformed and non-transformed cells within the tumour microenvironment. Achieving this requires incremental and meticulous investigations, free from biases that might overemphasize the significance of individual findings. However, this is just the first step toward developing therapeutic strategies targeting tumour acidosis. It is also essential to understand how acidosis, in conjunction with other pathophysiological conditions of the tumour microenvironment, impacts various aspects of carcinogenesis. The complexity of this challenge is underscored by the number of conditions, their unknown spatial severity, and the sheer number of possible combinations. We recently found that tumour-relevant concentrations of ammonia are cytotoxic to cancer cells^52^. Surprisingly, this cytotoxicity was highly dependent on the extracellular pH, suggesting that acidosis may modulate the effects of other metabolites in unexpected ways. Furthermore, hypoxia and acidosis are interrelated; studies have demonstrated that the cellular response to hypoxia is diminished when cells are simultaneously exposed to lactic acidosis^53^. These findings emphasize the need to study tumour microenvironmental factors not in isolation but as a dynamic, interconnected network. Only by unravelling these complex interactions can we identify actionable targets and design therapies that account for the multifactorial nature of cancer progression.

## Acknowledgments

The author acknowledges all members of the Cell Signalling and Targeted Therapy lab for insightful discussions. J.M. is supported by a grant from the Erna and Olav Aakres foundation. O.M. is supported by Norwegian Cancer Society (grant 198119-2019) and Northern Norway Regional Health Authority/Helse Nord RHF (project HNF1547-20).

## Author contributions

J.M. and M.D wrote the manuscript. M.D. performed the experiments. All authors contributed to the experimental planning and the manuscript editing.

## Materials and Methods

### Adhesion Assay

The adhesion assay was performed in 96-well plates. The plates were coated overnight with 100 µl of freshly prepared substrate of Laminin (20 µg/ml diluted in 0.1M NaHCO_3_), Fibronectin (50 µg/ml diluted in 0.1M NaHCO_3_), or Collagen (50 µg/ml diluted in 0.02M Acetic Acid) at 4-8 °C. Before use, the 96-well plates were rinsed twice with PBS and rinsed once with serum-free media containing 0.1% BSA.

The cells subjected to the assay were cultured in media with pH 7.5, media with pH 6.8, or in media containing TGF-β1 (Peprotech, #100-21, 0.5 µg/ml) for two days before the adhesion assay was performed. The adhesion assay was performed as follows: Growth media above the cells was saved, transferred into sterile containers, and stored in the incubator (5% CO_2_, 37°C). The cells were then washed once with PBS and trypsinized for 5 minutes. The trypsinized cells were then resuspended in 10 ml of acclimatized media with pH 7.5 (37°C) and centrifuged at 500g for 5 minutes. The supernatant was then removed and replaced with 1.1 ml of the saved growth media. The cells were counted using the DeNovix CellDrop Automated Cell Counter, and a final cell concentration of 0.4 x 10^6 cells per ml was prepared by adding the required amount of saved growth media. Next, 100 µl of this cell solution was dispensed into individual wells of a precoated 96-well plate. After 10-40 minutes in the in the CO_2_-incubator, the wells were washed thoroughly three times with 200 µl of PBS before 100 µl of PBS was added to each well. Thereafter, 100 µl of RT CellTiter solution was added to each well before the plate was placed on a stirrer for 20 minutes (speed: 220). The solution in each well was then resuspended, and 100 µl was transferred to a new white 96-well plate. Finally, luminescence was measured using the Spark® multimode microplate reader (shaking: double orbital, Luminescence: 1 sec.).

### Antibodies

The following primary antibodies were used in these studies: E-cadherin (Cell Signalling Technology, #14472), N-cadherin (Merck, #C3865-100µL), Paxillin (Cell Signalling Technology, #50195), and Phalloidin (Life Technologies; #A22287, Abcam; #ab235138 and #ab176753). The following secondary antibodies were used: HRP-conjugated goat anti-rabbit and anti-mouse (BD Pharmingen, #554021 and #554002). Alexa Fluor 568- and 488-conjugated antibodies were purchased from Life Technologies and used at a 1:1000 dilution.

### Cell Counting

Cell counting was performed using the DeNovix CellDrop Automated Cell Counter. A 100 µL aliquot of cells was mixed with 100 µL of PBS containing Acridine Orange (4.975 mg/L, Sigma #A6014). Subsequently, 40 µL of this mixture was applied to the cell counter and the number of cells was measured.

### Cell Culture

The studies were performed with five human carcinoma cell lines: lung cancer A549, epidermoid cancer A431, colorectal cancer HT-29, breast cancer MDA-MB-231, and colorectal cancer HCT116. All cell lines were obtained from the American Type Culture Collection (ATCC) and were regularly tested for mycoplasma contamination.

A549 cells were cultured in low-glucose Dulbecco’s Modified Eagle’s Medium (DMEM, Sigma, #D6046) supplemented with 9% FBS (Biowest, #S181B) and Penicillin-Streptomycin (90 U/ml and 0.9 mg/ml, Sigma-Aldrich, #P0781). In experiments where the pH was adjusted to pH 7.5, A549 cells were cultured in DMEM (Sigma, #D5030) supplemented with 9% FBS (Biowest, #S181B), Penicillin-Streptomycin (90 U/ml and 0.9 mg/ml, Sigma-Aldrich #P0781), GlutaMAX (Gibco, 0.9X, #35050-061), glucose (0.881 mg/ml, Sigma, #G8644), and sodium bicarbonate (1.11 g/L, Sigma, #S8761). In experiments where the pH was adjusted to 6.8, A549 cells were cultured in DMEM (Sigma, #D5030) supplemented with 9% FBS (Biowest, #S181B), Penicillin-Streptomycin (90 U/ml and 0.9 mg/ml, Sigma-Aldrich, #P0781), GlutaMAX (Gibco, 0.9X, #35050-061), glucose (0.872 mg/ml, Sigma, #G8644), and sodium bicarbonate (0.26 g/L, Sigma, #S8761).

A431 and MDA-MB-231 cells were cultured in high-glucose DMEM (Sigma, #D5796) supplemented with 9% FBS, Penicillin-Streptomycin (90 U/ml and 0.9 mg/ml, #P0781), and sodium pyruvate (0.9 mM, #S8636). In experiments where the media was adjusted to pH 7.5, A431 and MDA-MB-231 cells were cultured in DMEM (Sigma, #D5030) supplemented with 9% FBS (Biowest, #S181B), Penicillin-Streptomycin (90 U/ml and 0.9 mg/ml, #P0781), GlutaMAX (Gibco, 0.9X), glucose (0.835 mg/ml, Sigma #G8644), sodium pyruvate (0.9 mM), and sodium bicarbonate (1.41 g/L, Sigma #S8761). In experiments where the media was adjusted to pH 6.8, A431 and MDA-MB-231 cells were cultured in DMEM (Sigma, #D5030) supplemented with 9% FBS (Biowest, #S181B), Penicillin-Streptomycin (90 U/ml and 0.9 mg/ml, #P0781), GlutaMAX (Gibco, 0.9X), glucose (0.848 mg/ml, Sigma, #G8644), sodium pyruvate (0.9 mM), and sodium bicarbonate (0.25 g/L, Sigma, #S8761).

HT-29 and HCT116 cells were cultured in McCoyʹs 5A Medium (Sigma, #M4892) supplemented with 10% FBS (Biowest, #S181B) and Penicillin-Streptomycin (100 U/ml and 1 mg/ml, Sigma #P0781) with the pH adjusted to 7.5 by use of sodium bicarbonate (2.37 g/L, Sigma, #S8761). This medium was also used as control medium in experiments. In experiments where the pH was adjusted to 6.8, HT-29 and HCT116 cells were cultured in McCoyʹs 5A Medium (Sigma, #M4892) supplemented with 10% FBS (Biowest, #S181B), Penicillin-Streptomycin (100 U/ml and 1 mg/ml, Sigma-Aldrich, #P0781), and sodium bicarbonate (0.37 g/L, Sigma #S8761).

All cell lines were cultured in a HERACELL CO_2_ incubator in the presence of 5% CO_2_ at 37°C. For short-term experiments (up to one week), the media was changed every day. For long-term experiments the media was changed every second day. Before each media change, the media was acclimatized in the CO_2_ incubator for a minimum of 1 hour to reach the desired pH and temperature.

All cell lines were stored according to the supplier’s instructions and used within 6 months after resuscitation of frozen aliquots. At the start of each experiment, cells in culture flasks were washed with PBS and trypsinized before being counted and distributed into the desired vessels. Subsequently, the cells were allowed to grow overnight in DMEM medium (as described above). The next day, the medium was replaced with experimental media adjusted to pH 7.5 or 6.8, depending on the specific requirements of the experiment.

### Immunocytochemistry and Microscopy

Cells grown on round 12 mm coverslips were fixed for 10 minutes in cold (-20° C) methanol and rinsed twice with cold PBS. They were then incubated in 2% BSA in PBS-T for 15 minutes at room temperature to block nonspecific binding. Subsequent incubations with the indicated primary antibodies and fluorescence-conjugated secondary antibodies were performed in PBS-T (containing 1% BSA) at room temperature for 1 hour. Coverslips were rinsed four times for 2 minutes each with PBS-T following both incubations. Coverslips were subsequently stained with DAPI (0.25 ng/ml) and mounted using VECTASHIELD® Antifade Mounting Medium.

For microscopy, we used a DeltaVision Elite microscope (GE) running SoftWoRx 1.0 software utilizing either a 60x or 100x objective. Image channels were acquired sequentially using appropriate filter sets for DAPI, Alexa Fluor 568, and Alexa Fluor 488. Ordinary phase-contrast micrographs were obtained using an AX10 microscope (Zeiss) equipped with a Retiga 6000 monochrome camera. In all experiments, images shown in individual panels were acquired using identical exposure times or scan settings and were adjusted identically for brightness and contrast using Photoshop CS5 (Adobe).

### Immunofluorescence and Cell Shape Analysis

Cells were seeded onto coverslips, fixed in 4% paraformaldehyde and permeabilized with 0.2% Triton X-100. For F-actin staining, cells were incubated with either FITC- or TRITC coupled Phalloidin (#ab235138) or Phalloidin-iFluor 488 (#ab176753) and DAPI. Stained cells were imaged using an Olympus IX71 inverted microscope equipped with a Retiga SRV CCD camera and the accompanying Micro-Manager 2.0 beta software. Cell shape analysis was performed using ImageJ software.

### Long-Term Experiment

Cells were seeded at a specific density and cultured in 25 cm² flasks and 20 cm^2^ dishes with experimental media at pH 7.5 or pH 6.8. The cells were trypsinized, and protein lysates and samples for qPCR were collected once a week, for a period of six weeks. The same cells were also reseeded into new flasks after each collection.

### Viability with CellTiter-Glo

Cells were seeded into 96-well plates and cultured as indicated. Following treatments, the CellTiter-Glo Luminescent Cell Viability Assay (Promega) was used to measure ATP levels, as instructed by the manufacturer. Fluorescence measurements were obtained using a Spark® multimode microplate reader.

### Western Blot Analysis

Samples for Western blotting were harvested in 1X SDS buffer (50 mM Tris, pH 6.8, 2% SDS, and 10% glycerol) and boiled for 5 minutes. Protein concentrations were then measured using the BCA Protein Assay Kit (23227; Pierce), and lysates were calibrated. Bromophenol blue and DTT were added to final concentrations of 0.1% and 100 mM, respectively, before the samples were boiled and run on SDS-PAGE gels at 120/160V for 90 minutes. For blotting, a semi-dry transfer was performed at 100 mA per blot for 1 hour using a buffer containing 14 mM glycine, 48 mM Tris, 0.03% SDS, and 0.15% ethanol. Membranes were stained with Ponceau S before being blocked in 5% dry milk in 1X PBS-T for 30 minutes. Incubation with the primary antibody was carried out overnight at 4°C. Membranes were washed six times with 1X PBS-T before being incubated with the secondary antibody for 1 hour at room temperature. Membranes were then washed six times using 1X PBS-T. Finally, membranes were developed using the SuperSignal West Femto Chemiluminescent Substrate (Pierce, #34095) and the SuperSignal West Pico PLUS Chemiluminescent Substrate (Pierce, #34078) on Hyperfilm ECL (18 × 24; GE28-9068-37; GE Healthcare) in a Curix 60 (AGFA) or by chemiluminescence detection using the ImageQuant LAS 3000 (GE Healthcare).

### Wound Healing Assay

Cells were detached from the tissue culture dish and counted as described in the Adhesion Assay. Thereafter, the cells were seeded onto a 96-well ImageLock plate (Essen Bioscience #4379) and placed into the incubator (5% CO_2_ and 37°C). The next day, the culture media was replaced with fresh media at either pH 7.5 or 6.8, and the cells were allowed to grow for two days, reaching 100% confluency. Subsequently, the Incucyte® 96-Well WoundMaker Tool was used according to the manufactureŕs protocol. Before use, the WoundMaker tips were first washed in 45 ml of distilled water for 5 minutes and then in 45 ml 70% ethanol for 5 minutes. The tips were then used in a sterile cabinet to create wounds on the 96-well ImageLock plate (Essen Bioscience #4379). After use, the tips were washed in 45 ml 0.5% Alconox for 5 minutes, followeed by washing in 45 ml Virkon for 5 minutes. They were then washed in 45 ml of distilled water for 5 minutes and finally in 45 ml 70% Ethanol for 5 minutes. After the wounds were created, the cells were placed into an Incucyte incubator with 5% CO_2_ and 37°C, and the Incucyte® Scratch Wound Analysis was performed according to the manufacturer’s instructions.

### qPCR

After incubation under pH 7.5 or pH 6.8, cells were harvested, and total RNA was isolated using TRIzol™ Reagent (Thermo Fisher Scientific, Invitrogen, # 15596018) according to the manufacturer’s instructions. The mRNA expression of E-Cadherin (CDH1), N-Cadherin (CDH2), SNAIL1 (SNAI1), SNAIL2 (SNAI2), ZEB1, ZEB2 and TWIST1 was analysed by quantitative PCR. A total of 1 µg of RNA was subjected to reverse transcription using the High-Capacity cDNA Reverse Transcription Kit (#4374966) following the manufacturer’s protocol and analyzed by qPCR using the PowerUp™ SYBR™ Green Master Mix (Thermo Fisher Scientific, #A25918) according to the manufacturer’s instructions. Relative gene expression was calculated using the ΔΔCt method, with GAPDH as the reference gene. Specific primers used in this study are listed below.

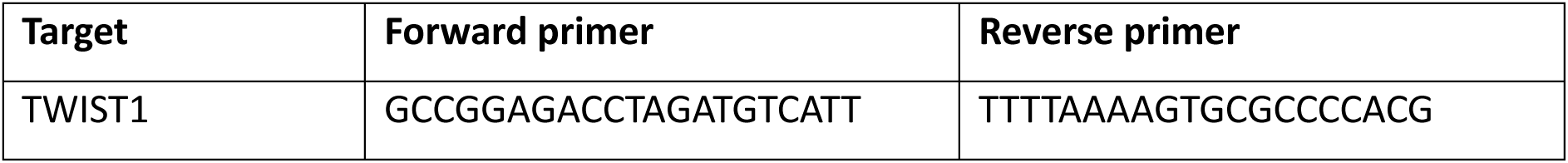
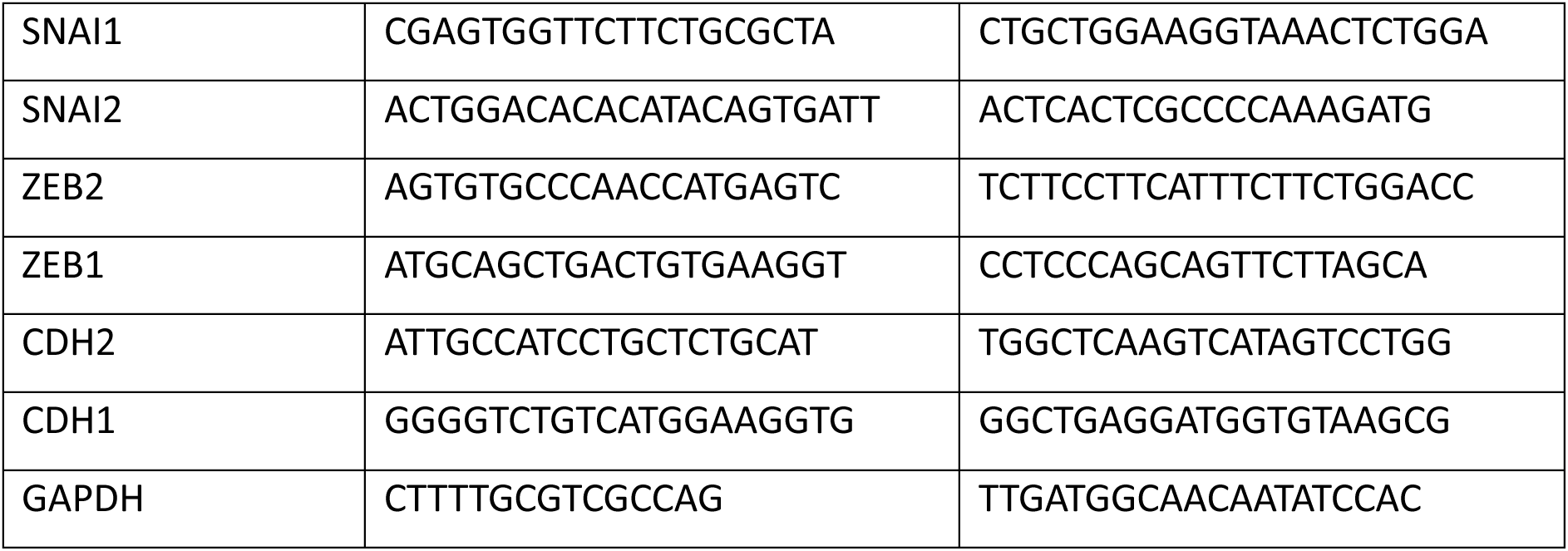

---

**Supplementary figure 1.**
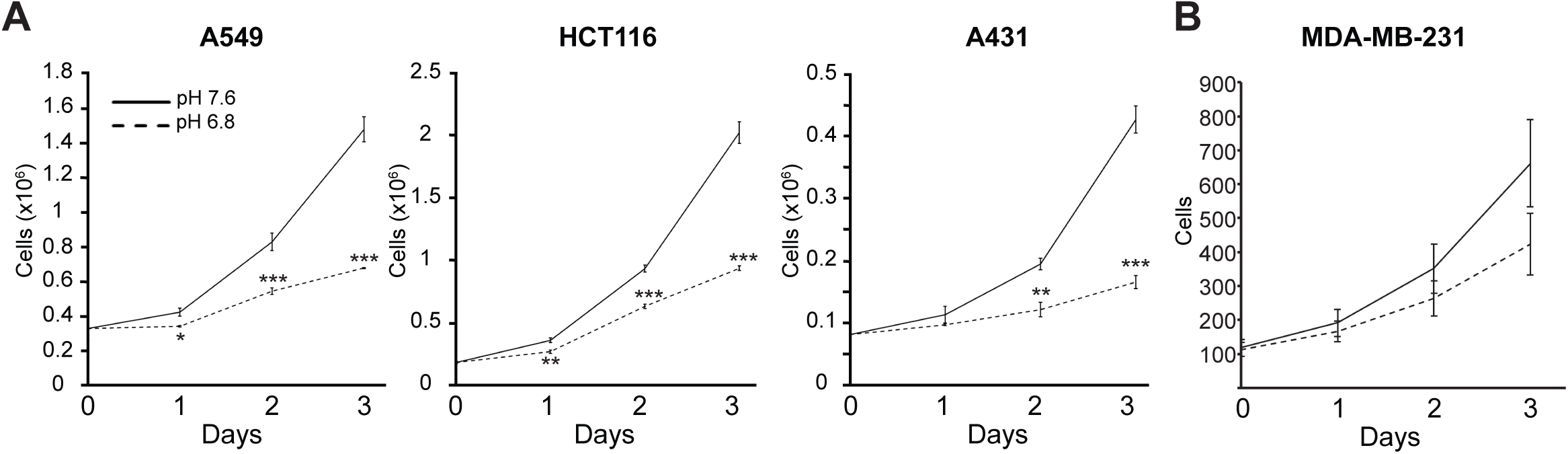
Low extracellular pH reduces the growth of cancer cells. **A**, The depicted cell lines were cultured for 3 days in media with either pH 7.5 or 6.8. Cells were counted daily using a DeNovix CellDrop Automated Cell Counter. Error bars represent the standard deviation of 3 technical replicates. Statistical p-values are based on a two-tailed Student’s t-test; *p<0.05, **p<0.01, ***p<0.001. **B**, MDA-MB-231 cells stably expressing nuclear mCherry were seeded on tissue culture plastic and cultured for three days while counted daily by Incucyte® Live-Cell Analysis System based on fluorescence. Source data are provided in the Source Data file.

**Supplementary figure 2.**
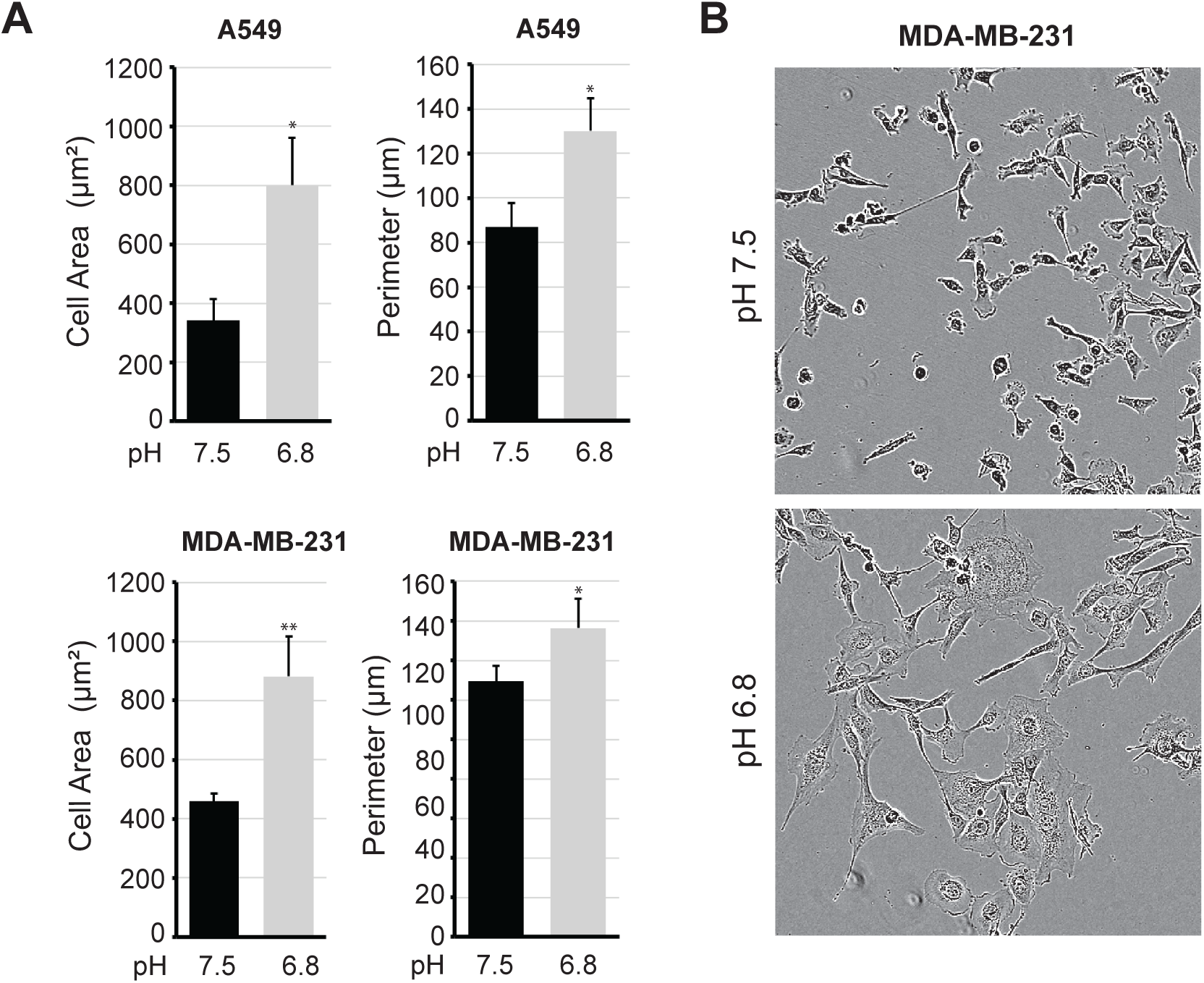
The morphology of MDA-MB-231 and A549 cells is affected upon acidosis. **A**, A549 and MDA-MB-231 cells were seeded on coverslips and cultured in growth media with either pH 7.5 or pH 6.8 for two days, before being fixed and stained with TRITC-phalloidin and DAPI. Imaged cells were subjected to cell shape analysis using ImageJ. Error bars represent the standard deviation based on three biological replicates with n>30. Statistical analysis was performed using two-tailed Student t-test; *p<0.05, **p<0.01. Source data are provided in the Source Data file. **B**, MDA-MB-231 cells were seeded on tissue culture plastic and cultured in growth media with either pH 7.5 or pH 6.8 for two days. Microscopic images were acquired by Incucyte® Live-Cell Analysis System.

**Supplementary figure 3.**
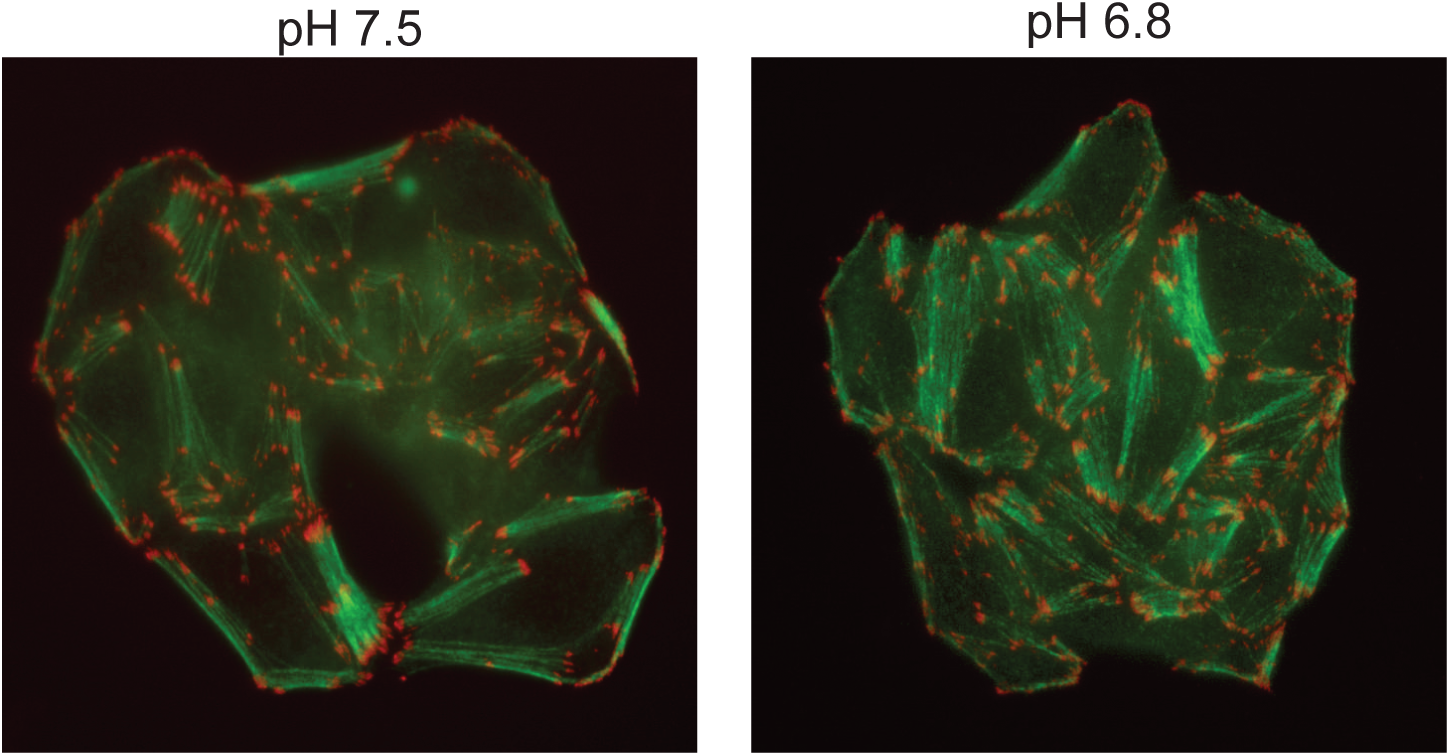
Immunofluorescence micrographs depicting F-actin (green) and Paxillin (red) in HT29 cells cultured for two days in media with either pH 7.5 or pH 6.8.

**Supplementary figure 4.**
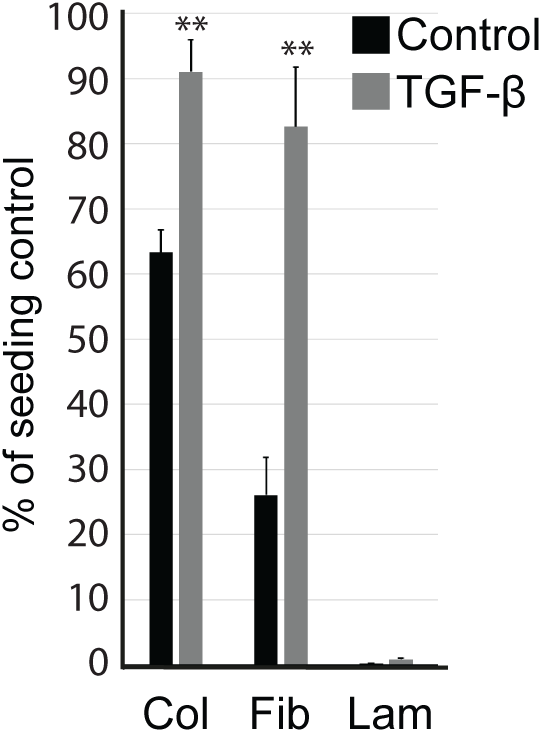
The relative adhesion to collagen (Col), fibronectin (Fib), and laminin (Lam) was assessed in A549 cells cultured for two days in either the absence or presence of TGF-β (5 ng/ml). Error bars represent the standard deviation of technical triplicates. The experiment was performed three times, yielding similar results. Statistical p-values are based on a two-tailed Student’s t-test; **p<0.01. Source data are provided in the Source Data file

**Supplementary figure 5.**
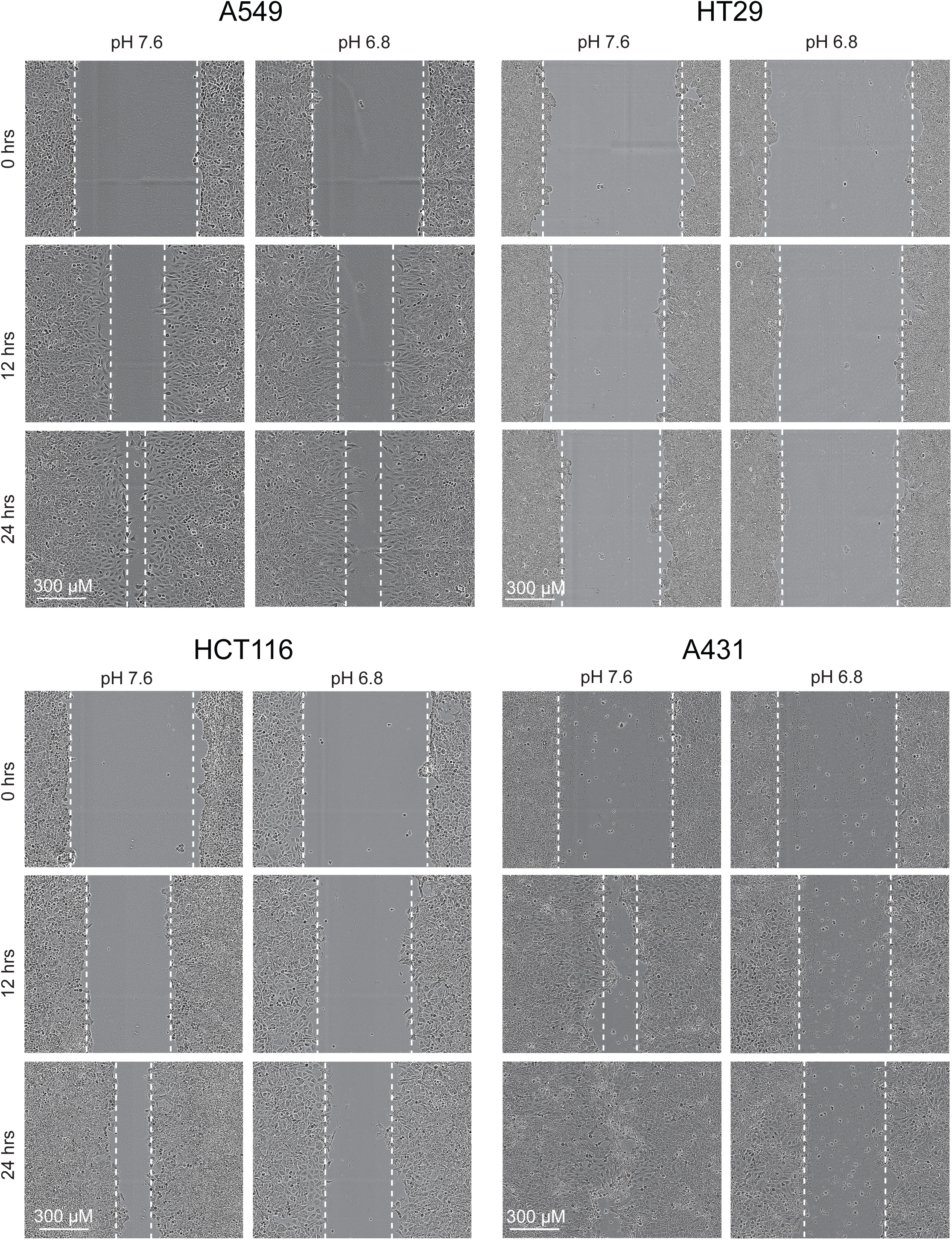
Phase-contrast microscopic images showing typical wound closure in the depicted cell lines. 0 hrs, 12 hrs and 24 hrs, indicate the time from when the wound was made.

**Supplementary figure 6.**
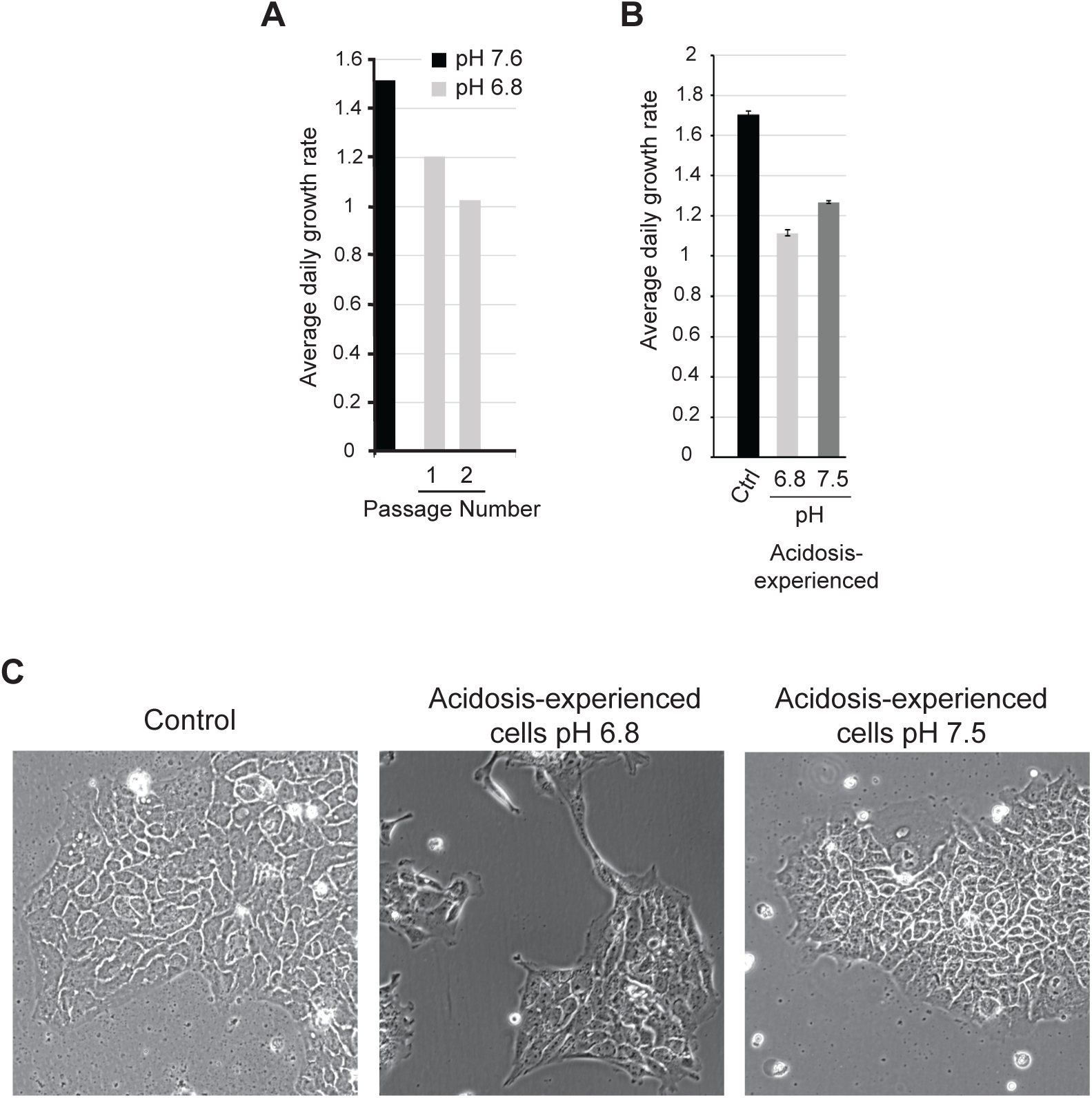
Prolonged exposure to low extracellular pH halts proliferation in A431 cells. **A**, A431 cells were seeded into media with pH 7.5. The following day, the media was changed to pH 6.8, and cells were allowed to grow under acidosis with regular media changes. After the second passage, A431 ceased to grow. The average daily growth rate is based on cell counting at each passage. The experiment was repeated three times, yielding similar results. Source data are provided in the Source Data file. **B**-**C**, After A431 cells had been cultured for 6 weeks in media with pH 6.8 (acidosis-experienced), they were split into two stocks and further cultured for 5 days in media with either pH 6.8 or 7.5). **B**, A column chart shows the average daily growth based on cell counting. Error bars depict the standard deviation of three biological triplicates. Source data are provided in the Source Data file. **C**, Images obtained by phase-contrast microscopy show cell morphology in the depicted samples.

**Supplementary figure 7.**
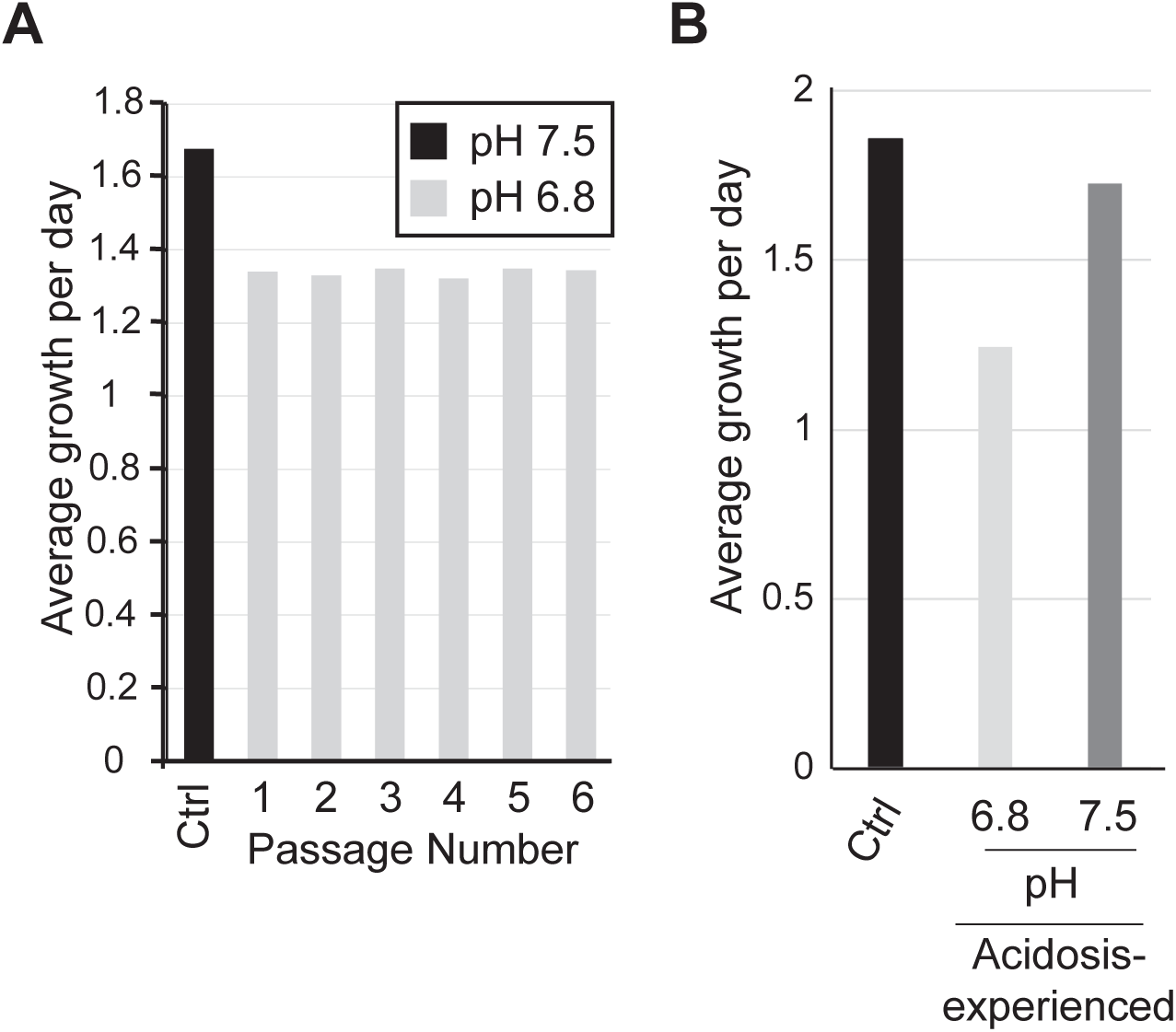
The proliferation of MDA-MB-231 cells is stably reduced during prolonged acidosis. MDA-MB-231 cells were cultured for 6 weeks in media with pH 6.8. **A**, Average daily growth rate is displayed for each consecutive passage following the start of the experiment. The experiment was repeated three times, yielding similar results. Source data are provided in the Source Data file. **B**, Cells cultured in media with pH 6.8 for four weeks (acidosis-experienced cells) were returned to media with pH 7.5, and cultured for one week before the average growth rate was determined. Source data are provided in the Source Data file.

**Supplementary figure 8.**
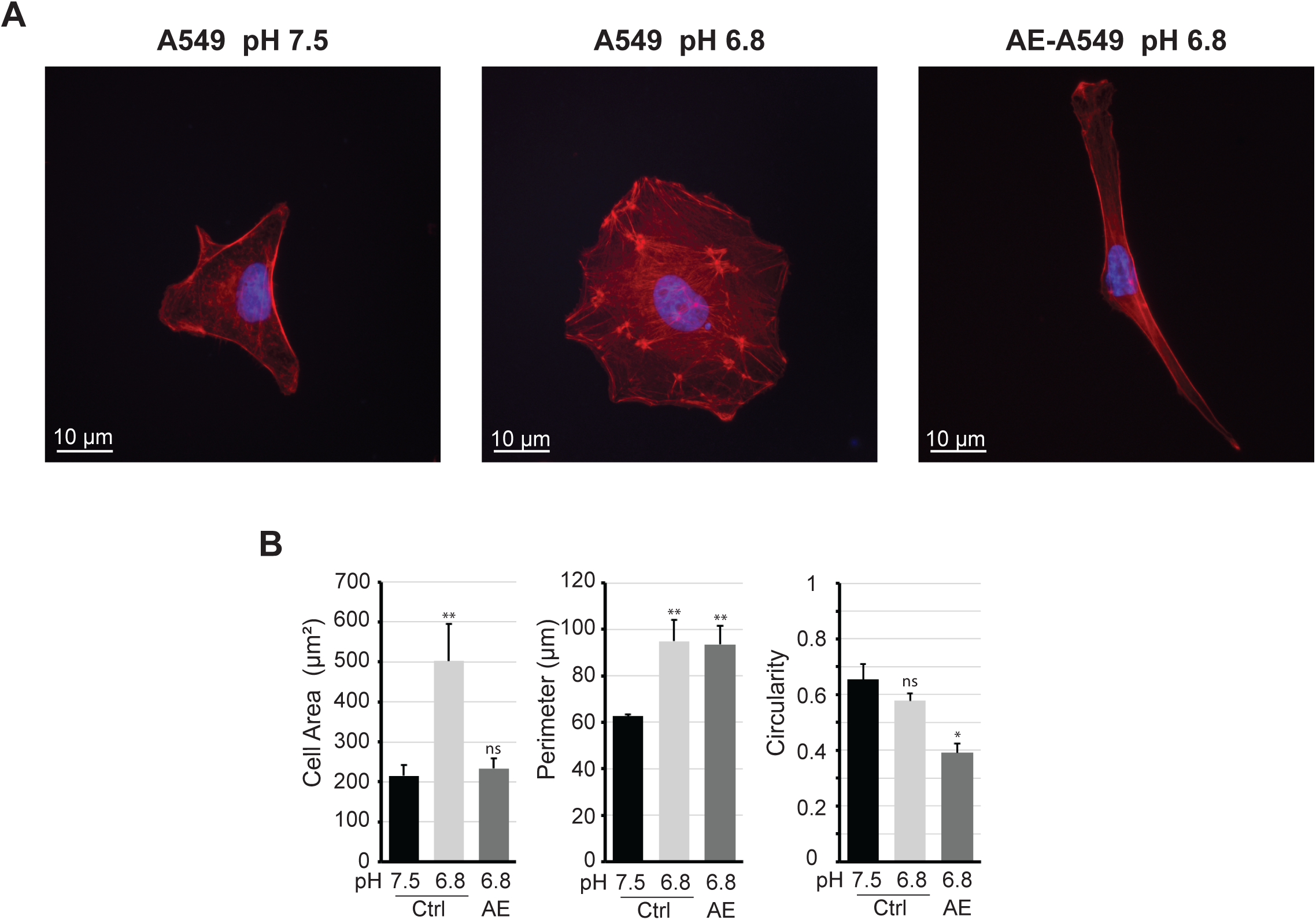
Acute acidosis and prolonged acidosis cause different morphological changes in A549 cells. **A**. Acidosis-experienced A549 (AE) and control A549 cells (Ctrl) were seeded on coverslips and cultured in growth media with the indicated pH for two days. The cells were then stained with TRITC-phalloidin and DAPI and imaged using immunofluorescence microscopy. The scale bar corresponds to 10 μm. **B**, Acidosis-experienced A549 (AE) and control A549 cells (Ctrl) cultured for two days were subjected to cell shape analysis using ImageJ. Error bars represent the standard deviation based on three biological replicates with n>30. Statistical analysis was performed using a two-tailed Student’s t-test; *p < 0.05, **p < 0.01. Source data are provided in the Source Data file.

**Supplementary figure 9.**
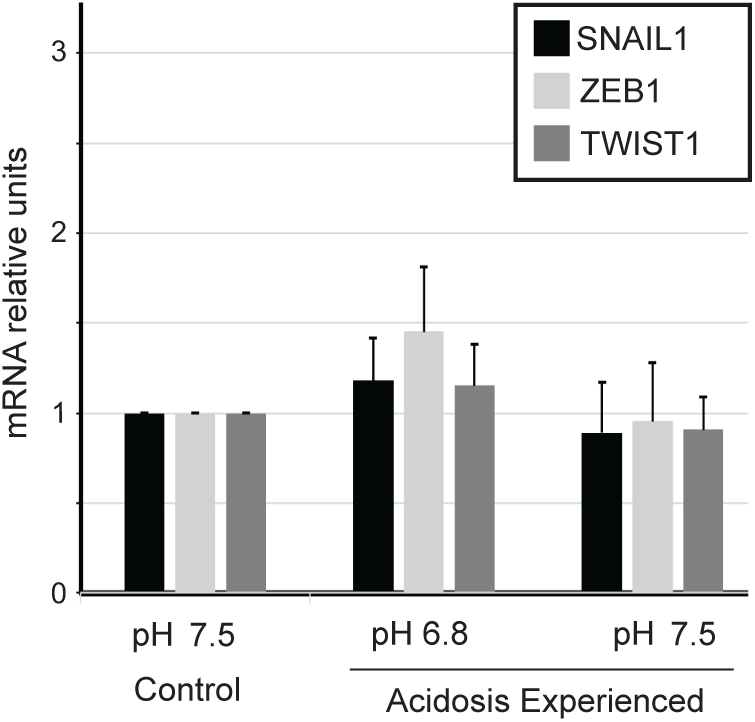
Prolonged acidosis does not influence the expression of SNAIL1, ZEB1 and TWIST1 in A549 cells. The expression of mRNA coding for SNAIL1, ZEB1 and TWIST1 was assessed by qPCR in A549 cells following the depicted treatments. “Acidosis-experienced” refers to A549 cells that had been cultured for more than 6 weeks in media with pH 6.8. Error bars depict the standard deviation of technical triplicates. Source data are provided in the Source Data file.

